# Single-cell transcriptomic analysis of HPV-related multiphenotypic sinonasal carcinoma uncovers MYB-HPV association

**DOI:** 10.1101/2025.04.21.649795

**Authors:** Avishai Wizel, Matthew D. A. Spence, Michael Mints, William Britton, Ogoegbunam Okolo, Victoria Yu, Isabel Cioffi, Salvatore Caruana, Scott H. Troob, William C. Faquin, Derrick T. Lin, Itay Tirosh, Sidharth V. Puram, Anuraag S. Parikh, Yotam Drier

## Abstract

Human papillomavirus (HPV)-related multiphenotypic sinonasal carcinoma (HMSC) is a rare tumor that morphologically resembles high grade adenoid cystic carcinoma (ACC), yet exhibits indolent clinical behavior. Both demonstrate *MYB* proto-oncogene upregulation, but HMSC lacks the *MYB* translocation typically seen in ACC. Transcriptional changes in HMSC tumors remain uncharacterized. We performed single-cell RNA sequencing (scRNA-seq) on a human HMSC tumor and compared expression profiles with published ACC and oropharyngeal squamous cell carcinoma (OPSCC) scRNA-seq datasets. Primary malignant cells from HMSC (n=134) and ACC (n=980) clustered separately, and HMSC lacked bicellular differentiation into luminal and myoepithelial cells, distinguishing it from ACC. A greater proportion of HMSC cells expressing HPV-related genes (HPVon) expressed MYB (83% vs. 62%, p=0.022) and MYB targets (p=6.4*10^-6^), suggesting an HPV-MYB association. This finding was validated in HPV-positive OPSCC, with 7/10 tumors showing MYB upregulation in HPVon versus HPVoff cells (p<0.05). A 264-gene signature from HPVon HMSC cells was also associated with worse prognosis in HPV+ OPSCC (p<0.003), suggesting an alternate role for HPV that has not been well characterized. Further validation of the HPV-MYB association and prognostically relevant HPV gene signature may improve patient stratification and therapeutic strategies in HPV-related malignancies.

## Introduction

Human papillomavirus (HPV)-related multiphenotypic sinonasal carcinoma (HMSC) represents a subset of HPV-related carcinomas that arise within the sinonasal tract and histologically resemble salivary adenoid cystic carcinoma (ACC). First described 10 years ago,^1^ HMSC remains poorly characterized, as it is an extremely rare tumor entity. HMSC exhibits a broad morphologic spectrum, reflecting features of both salivary-type and squamous cell carcinomas.^1–3^ Despite its typical high grade histology, HMSC has low potential for metastasis and is rarely fatal.^1^

Like ACC, myeloblastosis (MYB) genes are typically upregulated in HMSC.^4^ However, ACC harbors *MYB or MYBL1* chromosomal translocations in over 70% of cases,^5–7^ which juxtapose super-enhancers next to the MYB locus to drive expression.^5^ These translocations are notably absent in HMSC,^1^ thus suggesting an alternate mechanism of MYB upregulation. Moreover, MYB may facilitate paracrine interactions between myoepithelial and luminal cells in ACC, driving oncogenic NOTCH signaling in luminal cells.^5,8^ Conversely, HMSC lacks evidence of bicellular differentiation, suggesting MYB may act differently in these tumors.^2^

HMSC is also unique from ACC in harboring high-risk HPV33.^1–3^ HPV is known to drive oropharyngeal carcinogenesis through viral protein E6/E7-induced cell cycle entry and dysregulation of p53 and Rb. This leads to unchecked DNA damage, excessive DNA replication and genomic instability,^9^ but the predominant HPV subtypes driving these oropharyngeal cancers are HPV 16 and 18.^10^ Recent studies have also suggested the importance of HPV expression heterogeneity across malignant cells within HPV-positive (HPV+) oropharyngeal squamous cell carcinoma (OPSCC).^11^ Specifically, a subset of OPSCC tumor cells may lose HPV expression and enter a senescent phenotype, which drives therapy resistance, invasion, and worsened prognosis.^11^ In HMSC, though, the role of HPV and the impact of its potential expression heterogeneity remain unclear.

Despite their differences, distinguishing HMSC from ACC remains a diagnostic challenge,^3^ and some question whether this rare tumor is, in fact, a separate molecular entity. We employed single cell RNA-sequencing (scRNA-seq)^12,13^ to characterize transcriptional heterogeneity in HMSC with an eye towards understanding MYB expression and the role of HPV in these tumors, while providing a comparison with salivary ACC and HPV-positive oropharyngeal SCC. This work represents the first detailed transcriptional characterization of intra-tumoral heterogeneity in this rare disease entity.

## Results

### HMSC is genomically and transcriptionally distinct from ACC

To profile intratumoral heterogeneity in HMSC, we harvested a fresh tumor biopsy from a 56-year-old woman who underwent a left partial maxillectomy for recurrent poorly differentiated basaloid carcinoma. Pathology revealed solid sheets of cells with focal ductal differentiation **(Figure 1A)** and multifocal positivity for both P16 (**Figure 1B**) and MYB (**Figure 1C**). HPV PCR was positive for HPV type 33, supporting the diagnosis of HMSC and distinguishing the lesion from high grade, solid-type ACC. The freshly resected specimen was dissociated into a single-cell suspension (see **Methods**),^8^ flow-sorted to deplete nonviable and CD45-positive cells, and profiled by the SMART-seq2 protocol,^12^ which provided full-length transcripts for sequencing and analysis. Transcriptomes from a total of 155 HMSC single cells were retained after initial quality control (see **Methods)**.

**Figure 1.**
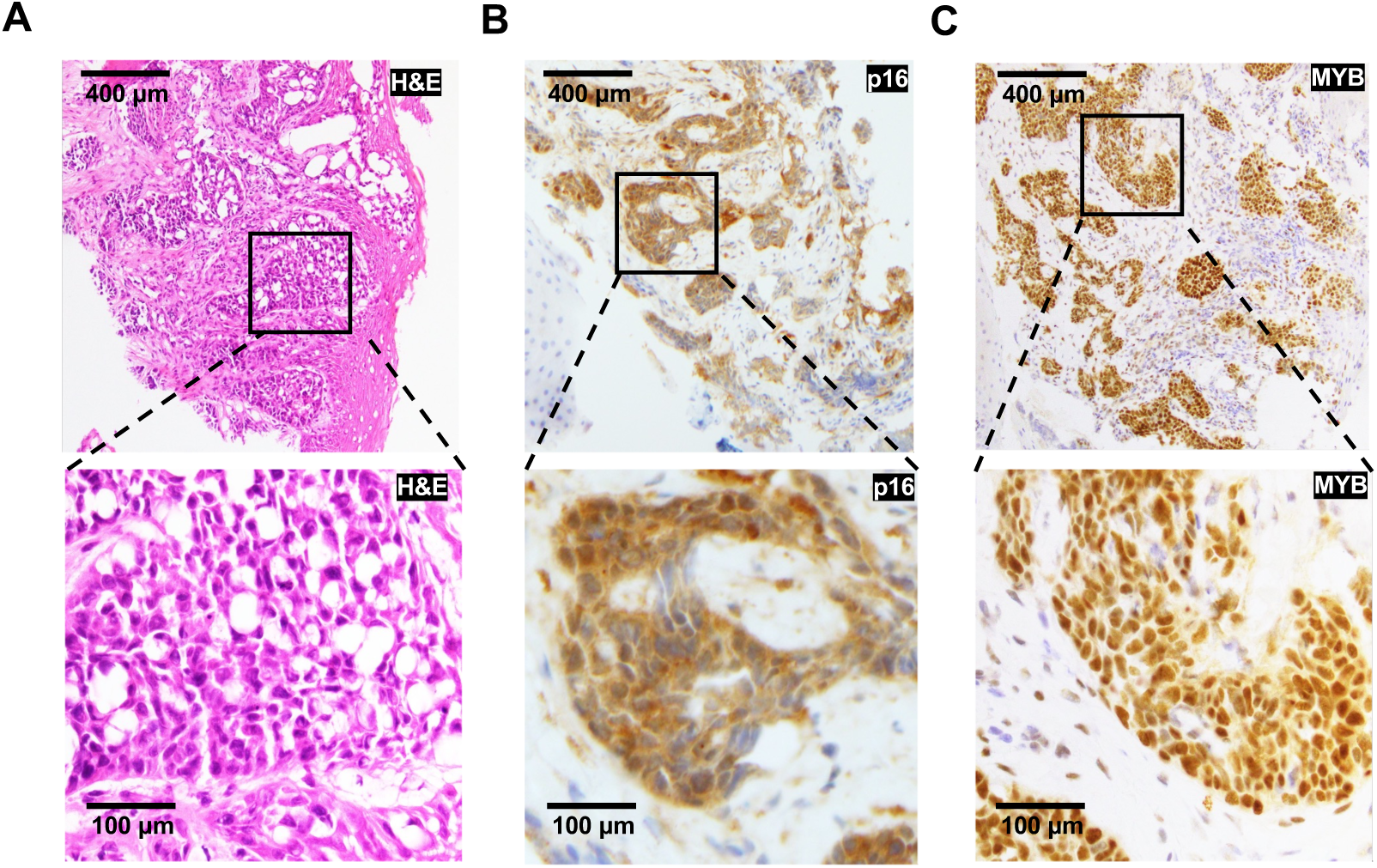
HMSC shows strong p16 and MYB staining. Histopathologic images of HMSC taken at 100X (top row, scale bar represents 400 µm) and 400X (bottom row, scale bar represents 100 µm) show (A) H&E staining, (B) strong and diffuse p16 immunohistochemical (IHC) staining, and (C) strong and diffuse MYB IHC staining.

We first sought to compare HMSC with ACC and combined our dataset with our previously published single cell transcriptomic analysis, which resulted in 980 primary malignant cells from seven head and neck ACC tumors after quality control.^8^ Expression analysis of known cell markers (see **Methods)** revealed that 143/155 cells in the HMSC dataset were malignant cells, with 134/143 cells that passed additional quality control and batch correction (see **Methods)**. Moreover, analysis of inferred copy number variations (CNV; see **Methods**) across the two datasets confirmed the distinction between malignant cells and non-malignant cells by demonstrating more CNVs for malignant cells. In addition, this analysis also revealed that malignant HMSC cells harbored a significantly higher degree of CNVs compared to malignant ACC cells (**Figure 2A, Supplementary Figure S1A**). Specifically, HMSC cells exhibited losses within chromosomes 4, 6, 9, 12-15, and 17, and gains within chromosomes 3, 18-20, and 22 (**Figure 2A**). HMSC cells clustered separately from ACC cells based on global gene expression (**Figure 2B, Supplementary Figures S1B-C**), supporting the hypothesis that these represent distinct cellular entities.

**Figure 2.**
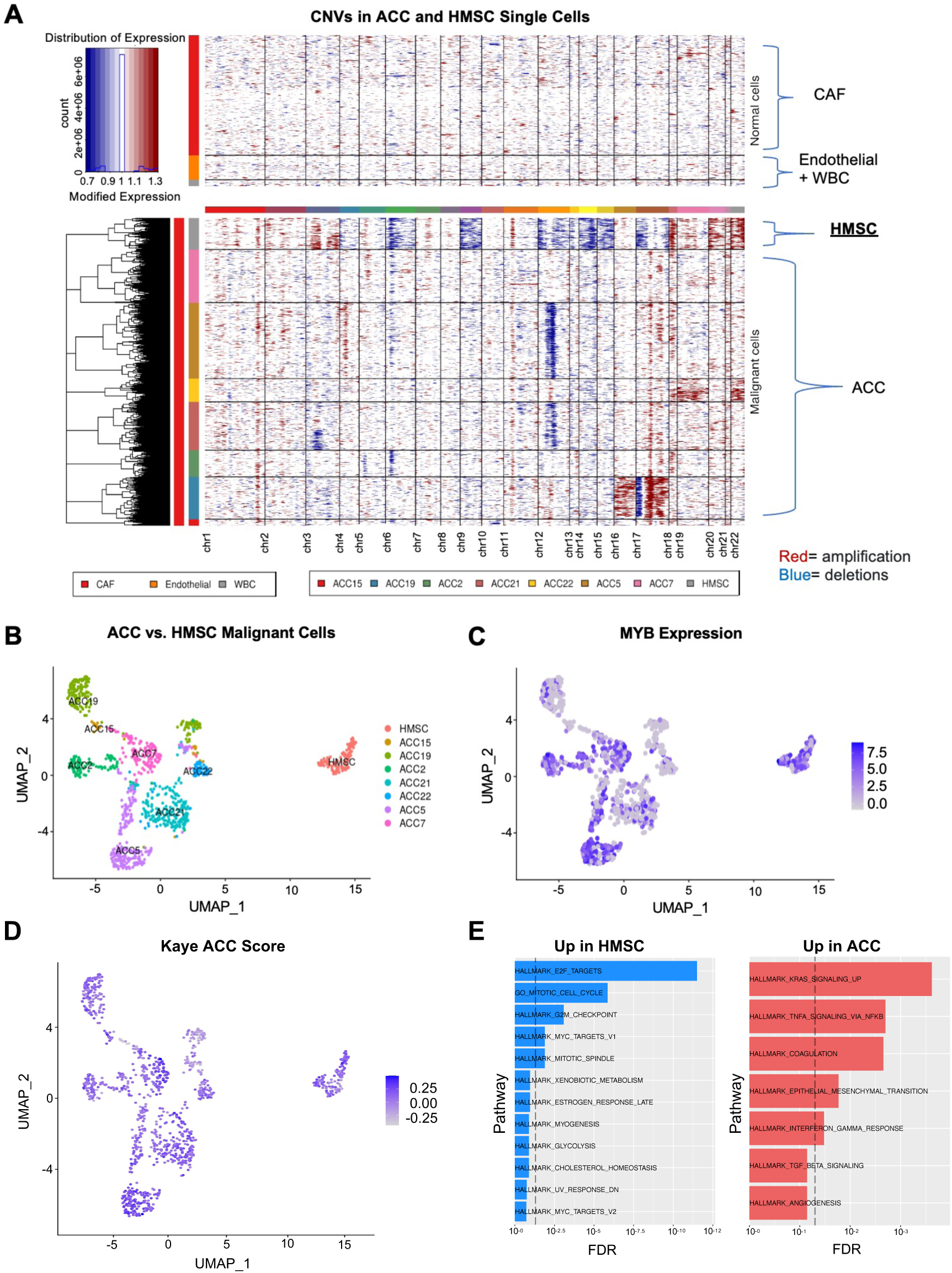
Single-cell RNA sequencing reveals that HMSC is genomically and transcriptionally distinct from ACC. A) Heatmaps show CNVs in HMSC and ACC non-malignant (top) and malignant (bottom) single cells. Columns represent chromosomal regions; red represents amplification, while blue represents deletion. B) Uniform manifold approximation and projection (UMAP) shows that primary malignant cells from HMSC cluster separately from those of ACC tumors. All cells are colored by patient. C) UMAP shows MYB expression in primary malignant cells from HMSC and ACC tumors. D) UMAP shows Kaye ACC score in primary malignant cells from HMSC and ACC tumors. E) Bar plots show gene set enrichment analysis (GSEA) for pathways upregulated in HMSC relative to ACC (left panel, blue bars, dashed line represents false discovery rate (FDR) <0.05) and pathways upregulated in ACC relative to HMSC (right panel, red bars, dashed line represents FDR<0.05).

We then scored cells based on MYB expression and found that malignant HMSC cells and ACC cells demonstrated high MYB expression **(Figure 2C)**, validating previous studies that demonstrated moderate to strong MYB staining in a majority of HMSC cases.^4^ Given the histologic similarities of these two entities, we also scored malignant cells for a previously defined ACC expression signature consisting of 1,160 mRNAs and 22 miRNAs^18^ to examine the degree to which HMSC resembled ACC at the transcriptional level **(Figure 2D)**. Both ACC and HMSC cells scored highly for this signature. Thus, despite differences in global clustering and CNV profile, malignant HMSC cells shared common transcriptional features with ACC cells.

Differential gene expression analysis comparing HMSC and ACC malignant cells revealed 94 genes that were more highly expressed in HMSC and 49 genes more highly expressed in ACC (FDR < 0.1, >1.25 fold-change, **Supplementary Figure S1D**). Gene set enrichment analysis revealed that genes with higher expression in HMSC were enriched for E2F targets (**Supplementary Figure S1E)**, consistent with well described changes that occur with HPV infection.^32^ Additionally, genes upregulated in HMSC demonstrated an enrichment of mitotic cell cycle genes and G2M checkpoint genes (**Supplementary Figure S1E**), while genes with lower expression in HMSC were enriched for epithelial-to-mesenchymal transition (EMT) genes (**Supplementary Figure S1E**). To estimate if this was due to a higher rate of cycling cells in HMSC, we filtered out cycling cells from both HMSC and ACC and repeated the analysis **(Figure 2E** and **Supplementary Figures S1F-S1G**). Even after the removal of cycling cells, genes higher in HMSC were still enriched in E2F targets and cell cycle genes (**Figure 2E**). For genes higher in ACC, filtering out cycling cells also revealed enrichment of KRAS signaling and EMT genes (**Figure 2E**).

### Intratumoral expression heterogeneity in HMSC

We next explored malignant cell expression heterogeneity in HMSC. We utilized non-negative matrix factorization (NMF, see **Methods)** to uncover three distinct expression programs within malignant cells **(Figures 3A-B**). Program one was enriched with cell cycle genes, program two with TNF-a signaling via NF-kB and apoptosis genes, and program three with estrogen response, coagulation, and complement related genes (**Supplementary Figure S2A)**. Most cells showed either a strong program one or program two signature, with only a few cells strongly expressing program three **(Figures 3A-B).**

**Figure 3.**
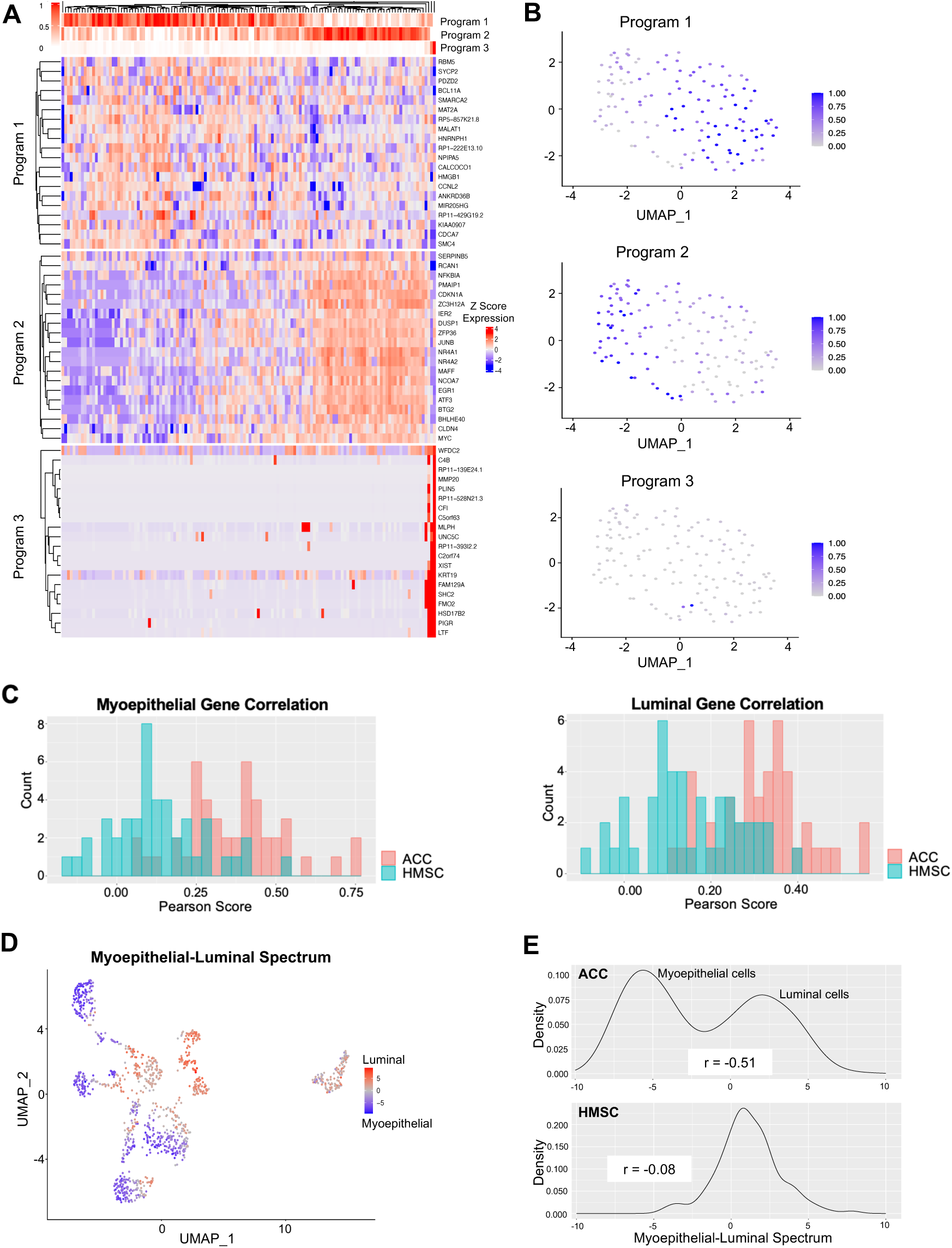
HMSC lacks bicellular differentiation. A) Heatmap shows top genes within three programs in HMSC. Cells are arranged with hierarchical clustering. B) UMAPs show expression of the three programs in malignant HMSC cells. C) Bar plots show counts of binned Pearson correlations among myoepithelial (left) and luminal (right) genes in ACC (red) and HMSC (blue). Correlations are generally weaker in HMSC than ACC. D) UMAP shows HMSC and ACC malignant cells on a spectrum from myoepithelial (blue) to luminal (red). HMSC cells demonstrate an intermediate phenotype. E) Line plots show smoothed densities of cells at points along the myoepithelial-luminal spectrum in ACC (top) and HMSC (bottom). ACC cells exhibit a bimodal distribution while HMSC cells exhibit an intermediate phenotype.

Notably, distinct myoepithelial and luminal expression programs were not uncovered in an unbiased fashion, as they were in ACC malignant cells.^8^ As a result, we next assessed the presence of distinct myoepithelial and luminal genes in HMSC. We annotated HMSC malignant cell clusters based on the expression of known myoepithelial **(Supplementary Figure S2B)** and luminal **(Supplementary Figure S2C)** markers.^8^ We then assessed the correlations within these myoepithelial and luminal gene sets for ACC and HMSC (**Figure 3C**). There were substantially higher correlations across luminal and myoepithelial genes in ACC than in HMSC, supporting the lack of distinct cell types in the latter. We then scored the HMSC and ACC malignant cells on a myoepithelial-luminal spectrum **(Figures 3D-E,** see **Methods)** and found that while ACC cells ranged from highly myoepithelial to highly luminal, HMSC cells largely scored toward the middle of the spectrum, suggesting that cells did not fall clearly into one or the other category. Thus, as previously described,^8^ ACC showed a bimodal, bicellular differentiation distribution; by contrast, HMSC cells displayed a single intermediate phenotype along this axis.

Given the prominent role of Notch signaling in ACC pathology,^33–35^ we also scored all malignant cells based on Notch signaling using known Notch target genes **(Supplementary Figure S3A)** and found that Notch was only activated in a handful of HMSC cells. We investigated the correlation between myoepithelial score and both Notch ligands and Notch targets in malignant HMSC cells (**Supplementary Figure S3B**). Unlike ACC, there was no significant correlation between the myoepithelial score and Notch ligand expression (r=-0.088, p=0.31), likely because myoepithelial score does not capture essential biological features in the context of HMSC. There was a weak but statistically significant positive correlation between myoepithelial score and Notch target expression (r=0.21, p=0.015), but this association was dependent on a single cell and was no longer significant once it was removed (r= -0.028, p=0.75). Taken together, these results suggest that Notch signaling is unlikely to play an essential role in HMSC oncogenesis and heterogeneity as it does in ACC.

### HPV33 is associated with MYB expression in HMSC

Given that HPV infection is a distinguishing trait of HMSC, we next characterized HPV expression in HMSC malignant cells. Plotting HMSC cells by number of reads per cell that map to the HPV33 genome revealed a bimodal distribution of HPV33 expression, with most cells expressing either 0 reads (henceforth HPVoff cells) or at least 24 reads (henceforth HPVon cells, see **Figure 4A)**. Most malignant cells (105/134, 78.4%) were HPVon (**Figures 4A-B**). HPVon cells were distributed similarly across different clusters and on the uniform manifold approximation and projection (UMAP) (**Figure 4B**), suggesting HPV status may not drive a primary axis of transcriptional variability across cells. 264 genes were differentially expressed between HPVon and HPVoff cells (FDR < 0.1, >1.25 fold-change), all of which were upregulated in HPVon cells (**Figure 4C, Supplementary Table 1)**. Moreover, genes higher in HPVon cells were enriched for E2F targets, suggesting HPV activates E2F factors in HMSC (**Figure 4D**).

**Figure 4.**
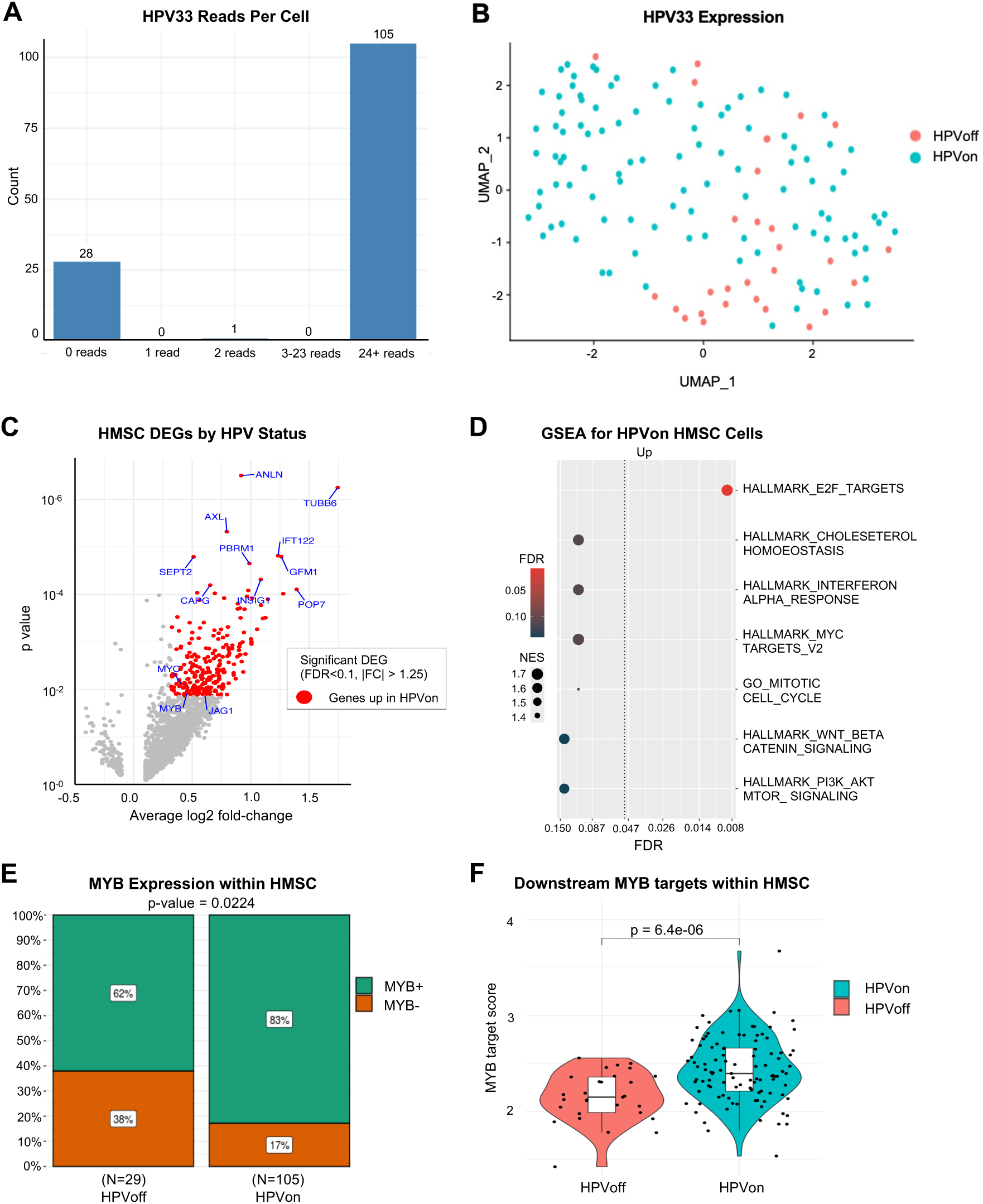
HPV is associated with MYB in HMSC. A) Bar plot shows counts of HMSC malignant cells with the specified numbers of HPV33 reads detected. Most cells had either 0 reads (28/134) or >24 reads (105/134). B) UMAP shows the distribution of HPVon cells across all HMSC malignant cells. HPVon cells are shown in blue. C) Volcano plot shows differentially expressed genes (DEGs) in HPVon HMSC malignant cells with upregulated genes shown in red (FDR<0.1, >1.25 fold-change). MYB is upregulated in HPVon cells. D) Dot plot shows GSEA for pathways that are upregulated in HPVon relative to HPVoff HMSC cells (dotted line marks FDR<0.05, color indicates significance, size indicates Normalized Enrichment Score (NES)). E) Stacked bar plots show proportion of MYB+ cells by HPV status. HPVon cells show greater MYB positivity than HPVoff cells (83% vs 62%, p=0.0224). F) Violin plot shows upregulation of downstream MYB targets in HPVon cells relative to HPVoff cells in HMSC (t-test, p=6.4x10^-6^).

One notable gene that was upregulated in HPVon cells was MYB (**Figure 4C**, fold-change=1.37, p<0.012, FDR<0.1). A significantly higher fraction of HPVon cells had overexpression of MYB (MYB+ cells) (**Figure 4E**, 83% vs. 62%, Fisher’s exact test p=0.022), and MYB targets were significantly upregulated in HPVon cells (**Figure 4F**, p=6.4 x 10^-6^). Taken together, these findings suggest HPV33 infection may drive MYB expression in HMSC tumors, instead of the canonical *MYB* translocations responsible for MYB upregulation in most ACC^5^. Given that MYB may drive alternate cell fates in ACC,^5^ including JAG1 and Notch1 expression in myoepithelial and luminal cells, respectively,^8^ we next explored expression of Notch ligands and targets in HPVon and HPVoff cells in HMSC **(Supplementary Figures S3C-F)**. While there was higher expression of JAG1 in HPVon cells (p=0.01), no other significant differences were observed in expression of Notch ligands or targets between cell types, consistent with the notion that Notch signaling may not play a significant role in HMSC oncogenesis.

### HPV-MYB association is also observed in HPV+ oropharyngeal SCC

To validate the observed HPV-MYB association in HMSC, we next assessed whether this association was retained in oropharyngeal squamous cell carcinoma (OPSCC), another HPV-related tumor of the head and neck. We reanalyzed scRNA-seq data from a previously published cohort of 10 patients with HPV+ OPSCC^11^ (**Supplementary Table 2**) to identify HPVon and HPVoff cells using similar approach. A significantly higher fraction of HPVon cells were MYB+ in 7/10 tumors (**Figure 5A**, Fisher’s exact test, p<0.05). Moreover, MYB targets were upregulated in HPVon cells (relative to HPVoff cells) in all 10 tumors assessed (**Figure 5B**), supporting the HPV-MYB association seen in HMSC. Finally, we evaluated MYB expression across all OPSCC tumors in The Cancer Genome Atlas (TCGA), finding significant upregulation of MYB in HPV+ tumors relative to HPV-tumors (**Figure 5C**).

**Figure 5.**
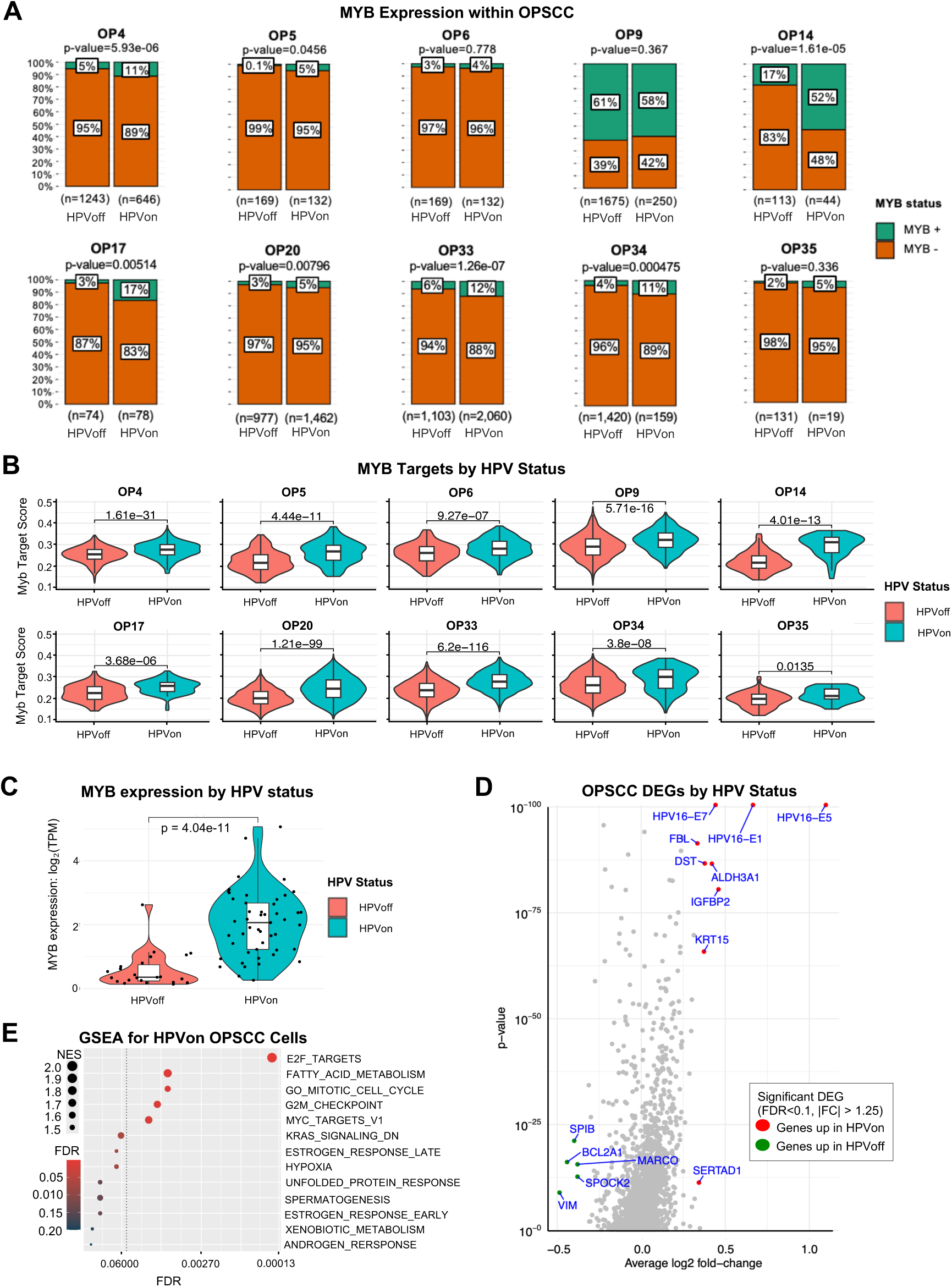
HPV is associated with MYB in OPSCC. A) Stacked bar plots show MYB+ cells by HPV status in OPSCC tumors. Higher MYB positivity (Fisher’s exact test, p<0.05) was seen in HPVon cells for 7/10 individual OPSCC tumors. B) Violin plots demonstrate upregulation of MYB targets in HPVon relative to HPVoff cells (t-test, p<0.05) in all 10 individual OPSCC tumors. C) Violin plot shows higher MYB expression in HPVon (right) versus HPVoff (left) OPSCC tumors from TCGA (t-test, p=4.04x10^-11^). D) Volcano plot shows DEGs in HPVon OPSCC malignant cells. Red represents genes upregulated in HPVon cells (FDR<0.1, >1.25 fold-change), and green represents genes upregulated in HPVoff cells (FDR<0.1, >1.25 fold-change). E) Dot plot shows GSEA for pathways upregulated in HPVon relative to HPVoff OPSCC cells (dotted line marks FDR<0.05, color indicates significance, size indicates NES).

We next sought to assess the degree to which HPV-related gene expression in these two tumor types was similar. In the cohort of HPV+ OPSCC, 14 genes were significantly differentially upregulated in HPVon cells (FDR<0.1, fold-change>1.25) (**Figure 5D, Supplementary Table 3**). These genes did not overlap with genes significantly upregulated in HMSC HPVon cells. However, as in HMSC, genes upregulated in HPVon cells were enriched for E2F targets in OPSCC as well (**Figure 5E**). We next devised a gene signature based on differentially expressed genes in HMSC and examined its significance in OPSCC tumors. All 10 tumors exhibited a statistically significant upregulation of this gene signature within their HPVon cells relative to the HPVoff cells (**Figure 6A)**, suggesting this signature may carry importance in OPSCC as well.

**Figure 6.**
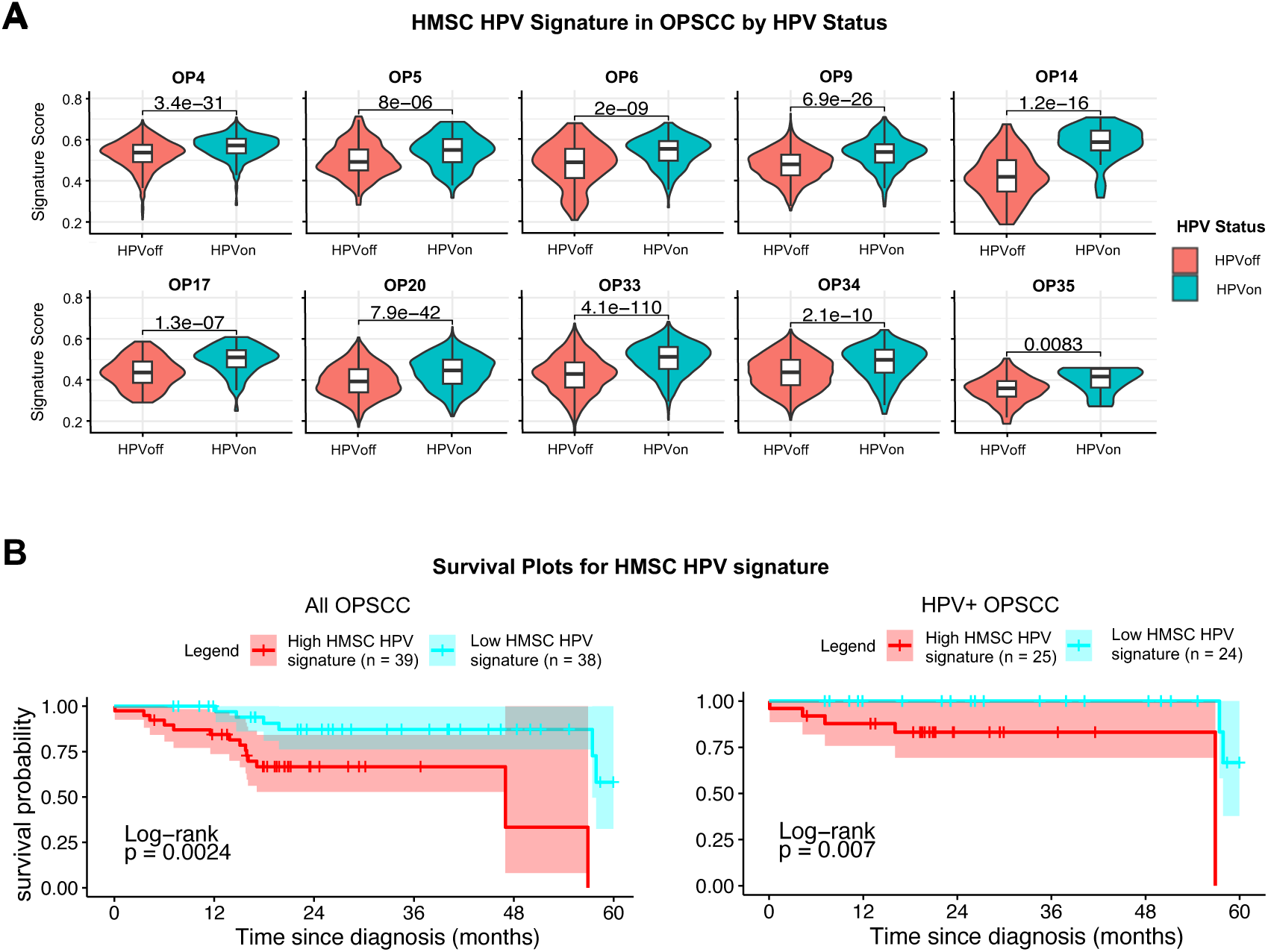
HMSC HPV signature is upregulated in HPV+ OPSCC and is associated with poor prognosis. A) Violin plots show upregulation of HMSC HPV signature in HPVon relative to HPVoff cells (t-test, p<0.05) for all ten individual OPSCC tumors (each p<0.01). B) Kaplan-Meier curves of TCGA all OPSCC (left) and HPV+ OPSCC (right) patients stratified by expression of HMSC HPV signature. Red survival curves represent patients with high expression of HMSC HPV genes and demonstrate poorer survival in both analyses (p<0.01).

To understand its prognostic significance, we examined survival associations of this gene signature in TCGA patients with OPSCC. While there was no survival association in patients with HPV-OPSCC (**Supplementary Figure S4A)**, we found a negative association between the signature and overall survival for all OPSCC patients as well as specifically amongst HPV+ OPSCC patients (**Figure 6B)**. Interestingly, there was no association between the signature and the stage of the tumor (**Supplementary Figures S4B-D**), suggesting the signature is an independent prognostic factor. As previously mentioned, some of the genes in the signature were associated with the cell cycle and E2F targets, so we repeated the analysis specifically excluding these genes. Similar associations were found (**Supplementary Figure S5**), suggesting these prognostic associations were not driven just by increased E2F activity and proliferation rate. In contrast with the notion that HPV expression in OPSCC is typically associated with a favorable prognosis,^11,36^ this signature may reflect an alternate, negative role of HPV in HPV+ OPSCC that is not well described and may contribute to variability in patient outcomes.^36,37^

## Discussion

HMSC represents an extremely rare entity, with less than 100 total cases described in the English literature. In this study, we present the first scRNA-seq analysis of this unique tumor type and provide a comparative analysis with scRNA-seq data from ACC and OPSCC. While our findings were limited by analysis of a single sample, they highlighted several important transcriptional differences between HMSC and ACC, including the distinct clustering of HMSC cells from ACC cells, the lack of bicellular differentiation in HMSC without defined myoepithelial or luminal programs, and the lack of oncogenic Notch signaling. These findings are consistent with previous histologic studies of HMSC^1–3^ and strongly support the notion that this is a distinct molecular entity, not to be characterized as a subset of ACC.

Notable expression differences between HMSC and ACC included lower expression of EMT genes and KRAS signaling in HMSC compared to ACC. EMT is hypothesized to be associated with invasion and metastasis in several epithelial tumor types,^38–40^ and in ACC we previously found upregulation of mesenchymal genes in myoepithelial cells.^8^ Decreased EMT in HMSC may contribute to the more indolent clinical behavior of these tumors, especially in comparison to their typical high grade histology at presentation.^3^ Downregulated KRAS signaling may have a similar impact, as activating mutations in the KRAS gene have been associated with poorer survival in head and neck squamous cell carcinoma.^41^ The most prominent non-cycling population of cells in this tumor was enriched for TNFa/NFkB signaling. Among its many effects, HPV infection has been shown to induce TNFa/NFkB signaling,^42^ which may have tumor suppressive activity.^43,44^ Upregulation in this pathway is associated with improved survival for patients with HPV+ tumors,^45^ again consistent with the slow-growing nature of HMSC tumors. These observations may begin to account for some of the distinct clinical behavior observed in HMSC compared to ACC and other similar histologic subtypes.

Despite these differences between HMSC and ACC, HMSC did display concordant expression of a previously described ACC transcriptomic signature,^18^ suggesting some expression similarities between these entities. These similarities may be related to upregulation of the MYB proto-oncogene in both tumor types,^4,7^ though by distinct mechanisms, as HMSC lacks the canonical *MYB* translocations described in ACC.^5–7^ Here, we uncovered an association between HPV infection and MYB expression that suggests an alternate, HPV-driven, mechanism of MYB upregulation in HMSC. Furthermore, this HPV-MYB association was validated in single cell and bulk transcriptional data in OPSCC, another HPV-related tumor type in the head and neck, which suggests it may be a pan-cancer association.

Interestingly, the downstream impacts of both MYB and HPV in HMSC may be different than those previously described in other tumor types. Specifically, regarding MYB, the lack of bicellular differentiation and Notch signaling or Notch ligand expression, as observed in ACC,^8^ suggests that MYB may have a distinct downstream role in HMSC. Further analysis of MYB binding sites in HMSC may help to elucidate these alternate mechanisms. In addition, regarding HPV, we uncovered 264 genes differentially upregulated in HPVon cells in HMSC. Although these genes were upregulated in HPVon OPSCC cells in all 10 tumors assessed, they were separate from genes differentially expressed by OPSCC HPVon cells in an unbiased analysis. Moreover, this gene signature was associated with poor survival in the TCGA OPSCC cohort, both when assessing all tumors and when filtering specifically to HPV+ tumors. In conjunction with data suggesting HPV positivity typically portends *improved* survival and outcomes in OPSCC,^11,36,37^ the poor survival association of this HPV-related signature suggests an alternate role of HPV that is less well characterized. Developing a better understanding of this signature may help to elucidate reasons for treatment failure in HPV+ OPSCC, a tumor that typically responds well to radiation therapy.^46^ Further investigation of these genes and this program may be instrumental in identifying future opportunities for new therapeutics for HPV+ OPSCC.

A major limitation of this study is the inclusion of only a single patient sample and limited number of cells. However, HMSC is an exceedingly rare tumor entity, and both the HPV-MYB association and the significance of the HMSC HPV gene signature were validated in OPSCC, suggesting these findings are robust. In addition, prospectively capturing more HMSC samples is often complicated by the fact that these tumors are often not identified until a definitive diagnosis is made on the final pathology report. Still, we acknowledge that the identified HMSC HPV gene signature requires further validation in larger patient cohorts and across diverse HPV+ tumor types to better understand its biological relevance, prognostic implications, and value as a therapeutic target. As the analyses presented are associational in nature, there is also the need for additional work to understand the mechanistic links between HPV and MYB expression. As a rare disease, HMSC currently lacks cell line or PDX models to validate these findings, underscoring the need for further model development in this rare tumor type.

In conclusion, this study reports the first scRNA-seq analysis of a rare HMSC tumor and highlights the transcriptional differences between HMSC and ACC. Whereas ACC cells typically exhibit either myoepithelial or luminal differentiation, HMSC cells demonstrated a single cellular phenotype. Our findings also highlighted a potential HPV-related mechanism of MYB upregulation within HMSC, distinguishing it from the *MYB-NFIB* fusion-driven overexpression in ACC. Further, we identified HPV-related genes in HMSC that were distinct from an HPV-related signature in OPSCC, suggesting alternate effects of HPV in these two tumor types. These HMSC HPV-related genes were associated with poor overall survival in TCGA OPSCC tumors, suggesting this alternate HPV effect may be relevant across HPV-related tumors. Further work is warranted to explore the biologic and prognostic utility of this program across HPV-related malignancies.

## Methods

### Human tumor sample

This study received approval from the Massachusetts Eye and Ear Institutional Review Board IRB, protocol 14-200H, in accordance with the Declaration of Helsinki. Prior to surgery, a patient at Massachusetts Eye and Ear with HMSC provided informed consent to participate in this project. A fresh tumor biopsy was obtained during the procedure, and the diagnosis of HMSC was confirmed by pathology.

### Tumor dissociation, cell sorting, and SMART-Seq2

Tumor dissociation, cell sorting, library preparation, and sequencing were performed as previously described.^8^ Briefly, the fresh tumor sample was mechanically and enzymatically dissociated to a single cell suspension using a Human Tumor Dissociation Kit (Miltenyi Biotec, Bergisch Gladbach, Germany). The cell suspension was filtered and resuspended, and cell viability was confirmed to be > 90% using trypan blue. Before fluorescence-activated cell sorting (FACS), cells were stained with 1 μM calcein AM (ThermoFisher Scientific, Waltham, MA, USA) and 0.22 μM TO-PRO-3 iodide (ThermoFisher Scientific) for viability and CD45-vioblue (Miltenyi Biotec) for immune cell depletion. Viable, singlet, CD45-cells were FACS sorted using 488 nm (calcein AM, 530/30 filter), 640 nm (TO-PRO-3, 670/14 filter), and 405 nm (Vioblue, 450/50 filter) lasers, standard forward scatter height versus area criteria, and calcein^high^/TO-PRO^low^ gates. Cells were sorted into 96-well plates containing TCL-buffer (QIAGEN, Hilden, Germany) with 1% β-mercaptoethanol and snap frozen prior to library prep.

Full length cDNA libraries were generated using a modified SMART-seq2 protocol.^12–14^ RNA purification was performed with Agencourt RNAClean XP beads (Beckman Coulter, Brea, CA, USA), reverse transcription with Superscript II or Maxima reverse transcriptase (ThermoFisher Scientific), and whole transcriptome amplification using KAPA HiFi HotStart ReadyMix (KAPA Biosystems, Wilmington, MA, USA). Tagmentation of cDNA libraries was performed with the Nextera XT Library Prep Kit (Illumina, San Diego, CA, USA) and sequencing was conducted as paired-end 38-base reads on a NextSeq 500 (Illumina).

### Processing of HMSC and ACC data and removal of non-malignant cells

HMSC data were aligned to the GRCh38.d1.vd1 reference with STAR version 2.5.2^15^ and counted with featureCounts.^16^ scRNA-seq data of ACC cells were taken from Parikh et al.^8^ Cells with less than 2,000 transcripts with at least 1 read, or more than 40% mitochondrial reads were removed from the analysis. Genes that were expressed in less than 3 cells across the combined HMSC-ACC cohort were also removed. Log_2_(TPM+1) values were calculated and used for downstream analysis. We then used the Seurat 4.1.0^17^ pipeline to perform dimensional reduction and clustering. We first scaled the data using the ScaleData function with the 15,000 most variable genes across the cohort, regressing out the number of detected reads and the percent of mitochondrial reads. Principal component analysis (PCA) was calculated with the same genes, and uniform manifold approximation and projection UMAP was done using the first 30 principal components (PC). Cells were grouped using the Louvain algorithm in Seurat, considering the first 30 principal components and setting the resolution to 1.

Each cluster was classified as malignant or non-malignant based on ACC score, defined as mean expression of known ACC markers,^18^ MYB expression, and expression of non-malignant markers -COL3A1 and COL1A2 as CAFs markers; CD79A and PTPRC as white blood cells markers; and VWF and PECAM1 as endothelial markers. All cells in the non-malignant clusters were removed from further analysis.

### Quality control and batch effect correction of HMSC data

To remove potential artifacts, we focused only on HMSC malignant cells with at least 2,500 transcripts. Potential batch effects between different plates were corrected with Seurat integration: 1,000 integration features were selected based on the 1,000 most variable features using the SelectIntegrationFeatures function. Integration anchors were found using the FindIntegrationAnchors function with k.filter = 50, and data was integrated using the IntegrateData function considering the 50 nearest neighbors,using k.weight = 50. Integrated data from HMSC malignant cells were then rescaled using the ScaleSeurat function with default arguments, and dimensionality reduction and clustering were performed using the Seurat pipeline, considering the first 10 principal components and a resolution of 0.5.

### Scoring for canonical pathways in HMSC and ACC

Mean expression of the genes JAG1, JAG2, and DLL1 were used for the NOTCH ligands score. For Myoepithelial score, Luminal score, and Notch targets score, we used the mean expression of genes as used in Parikh et al.^8^

We used MYB targets as identified in Drier et al.^5^ A cell was classified as MYB+ if MYB expression was > 1 TPM. An HMSC cell was classified as HPV+ if the number of reads mapped to the HPV33 genome was greater than 10. Cell cycle score was calculated using the Z-score mean of genes in GO mitotic cell cycle (GO:0000278). A cell was considered cycling if it had a cell cycle score of more than 0.3.

### Analysis of HMSC heterogeneity

cNMF (Consensus Non-negative Matrix Factorization on single-cell RNA-Seq data)^19^ version 1.5.0 python package with python 3.8.15 was used to determine expression programs from HMSC log_2_(TPM+1) values. We used the preprocess_for_cnmf function for batch correction specifying the plate variable under the “harmony_vars” parameter. We also set “max_scaled_thresh = 50” and “theta = 2”. Then we ran cNMF for 3 to 10 programs, and 20 iterations per run.

For differential analysis between HPVon and HPVoff HMSC cells, we used log_2_(TPM+1) values of the 3,000 most highly expressed genes in HMSC (of the 15,000 most variable genes in HMSC cells) and corrected for plate-based batch effects using linear regression with the argument test.use = “LR”.

Gene set enrichment analysis (GSEA) was calculated with the fgsea 1.20.0 R package,^20^ and visualized using hypeR version 2.0.1.^21^ Gene symbols with average log2 fold change, sorted in decreasing order, were used as inputs for GSEA. MSigDB Hallmarks V7.0^22^ and GO mitotic cell cycle (GO:0000278) were used in the enrichment analysis.

### Comparison between HMSC and ACC

The corrected HMSC expression data was then combined with the primary ACC cells with log_2_(TPM+1) values above. The combined HMSC-ACC data was rescaled with Seurat ScaleData regressing out the number of detected reads and percent of mitochondrial reads, using the 15,000 most variable genes of the combined dataset. PCA was calculated using the same number of variable genes and UMAP was calculated using the first 20 PC. To identify distinct cell populations within the dataset, we performed a clustering analysis using the Louvain algorithm with a resolution of 0.01.

For differential expression analysis, we initially used the log_2_(TPM+1) values of the 15,000 most variable genes across the combined dataset as an input to FindMarkers. Subsequently, we identified cycling cells as described above, removed them and then re-calculated the top 15,000 most variable genes. Log_2_(TPM+1) values of these genes across combined dataset of non-cycling cells were used as the input for FindMarkers, with the same parameters as before filtering.

### Copy number variation estimation of HMSC and ACC

Infercnv version 1.10.1^23^ was used to estimate copy number variation from integrated normalized expression values of the combined HMSC–ACC (primary and non-primary) cohort. Since the data were already normalized, step 3 (normalization by sequencing depth) and step 4 (log transformation of data) were skipped. Denoising was done with default Infercnv settings. Copy number aneuploidy score was defined as the square root of the average of deviation of diploidy squared.

### OPSCC single cell analysis

Raw data was downloaded from Gene Expression Omnibus with GEO accession GSE182227. Cells with less than 1,000 expressed genes and genes that were expressed in less than 3 cells were filtered out. Data was normalized using the Seurat NormalizeData function with log normalization and a scale factor of 10,000. For downstream analysis, we only used HPV+ tumors. A cell was identified as HPVon if it had a count of 3 or more for any HPV16 gene.

For differential analysis between HPVon and HPVoff OPSCC cells, we used the Seurat pipeline with the 5,000 most highly expressed genes (of the 15,000 most variable genes) and log2(TPM+1) values were used along with linear regression for patient identity to account for inter-patient variability.

### Statistical analysis and data visualization

Data analysis was performed in R (version 4.4.1)^24^ and ggplot2 version 3.5.1^25^ was used to generate figures. To generate heatmaps, we used ComplexHeatmap version 2.10.0,^26^ and we generated stacked barplots with the ggbarstats function from ggstatsplot package, version 0.12.5.^27^

We calculated Fisher tests using fisher_test and T-tests with t_test functions from the rstatix 0.7.0 R package^28^ with default arguments. False discovery rate (FDR) was calculated using the p.adjust function in stats package.

### TCGA data analysis

Transcriptomic and clinical data were downloaded using TCGAbiolinks^29^ with Data Release 42.0, while patients’ follow-up data were downloaded from the GDC portal. The follow-up data was restricted to a follow-up period of up to 60 months. RNA-seq data was log2 normalized and only samples originating from one of the following were included in the analysis: “Base of tongue, NOS”, “Oropharynx, NOS”, “Posterior wall of oropharynx”, “Tonsil, NOS”. HPV status was taken from cBioPortal^30^ and the TCGA pan cancer atlas 2018.^31^ For survival analyses, we split patients by median expression, and the signature was calculated with the mean of log2(TPM+1). Analysis and visualization were done using the TCGAanalyze_survival function from TCGAbiolinks. Creating the signature without cell cycle genes was done by excluding genes that were annotated as GO mitotic cell cycle (GO:0000278).

## Acknowledgements

Funding: Supported by V Foundation (S.V.P.), Cancer Research Foundation (S.V.P.), Barnes Jewish Hospital Foundation (S.V.P.), NCI 1K08CA237732 (S.V.P), the European Research Council Horizon 2020 grant 949029 (Y.D.), the Alon Fellowship of the Israeli Council for Higher Education (Y.D.), and NIDCR 1K08DE033093 (A.S.P.). The funding sources had no involvement in the design, conduct, and reporting of the research.

## Author Contributions

AW, MDAS, ASP, and YD were responsible for designing and executing the study as well as writing/editing the manuscript. DTL and SVP provided patient samples and edited the manuscript. MM, WB, OO, VY, IC, SC, SHT, WCF, and IT contributed to data interpretation, data analysis, and editing the manuscript. ASP, YD, SVP, and IT were also responsible for supervision.

