## Supplementary Figures for "Single-cell transcriptomic analysis of HPV-related multiphenotypic sinonasal carcinoma uncovers MYB-HPV association"

### Supplementary Figure 1

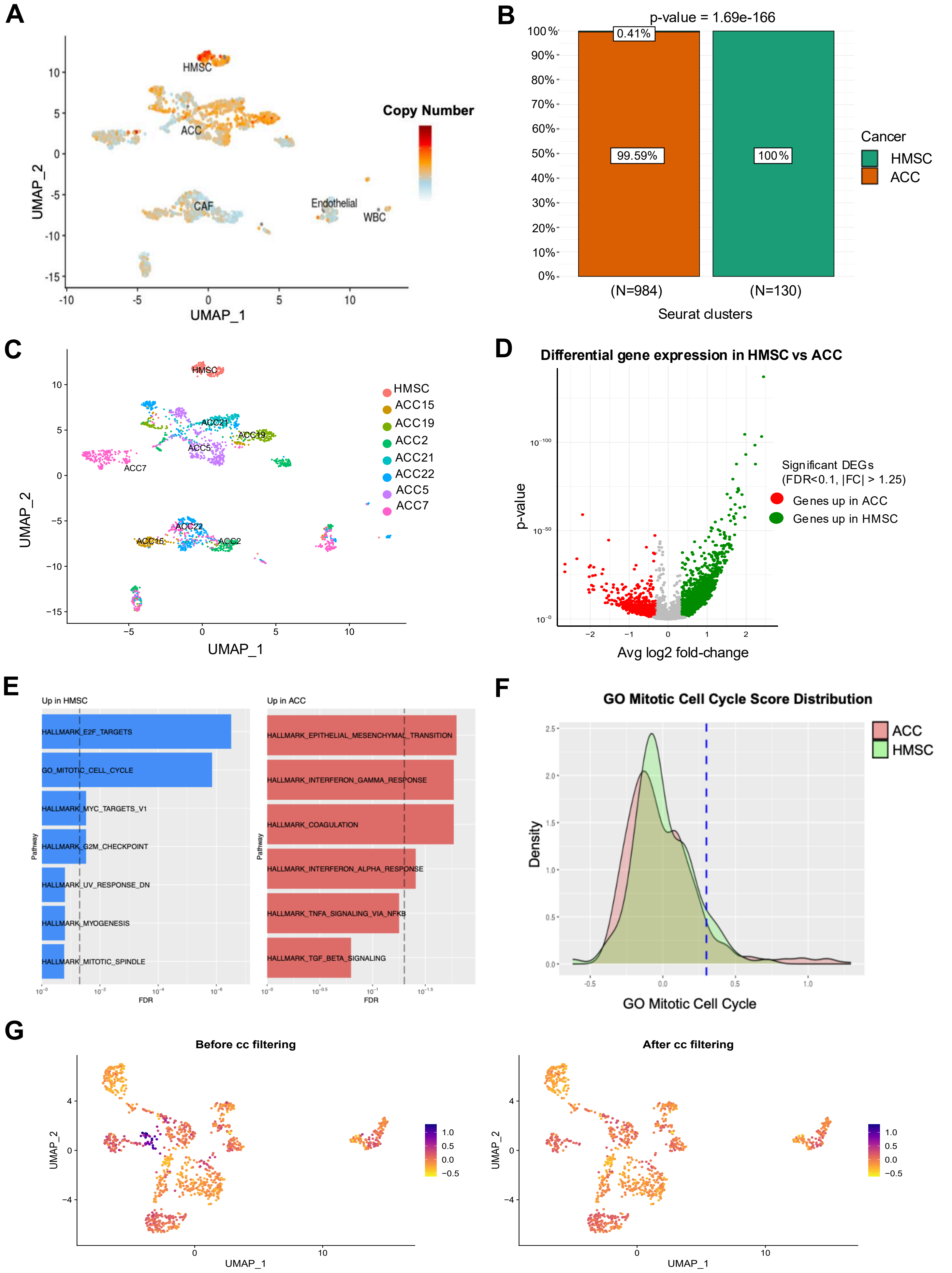

**Supplementary Figure 1. Additional classification of ACC and HMSC. Related to Figure 2.**

A. UMAP clustering of all cells, colored based on copy number variations (CNVs).
B. Stacked bar plot shows that unsupervised clustering separates HMSC (green) and ACC (orange) cells ($p<2x10^{-166}$).
C. Unsupervised UMAP clustering of all cells, colored based on tumor origin.
D. Volcano plot shows DEGs in HMSC relative to ACC. Green represents genes upregulated in HMSC cells (FDR<0.1, >1.25 fold-change) and red represents genes upregulated in ACC cells (FDR<0.1, >1.25 fold-change).
E. Bar plots show gene set enrichment analysis (GSEA) for pathways upregulated in HMSC relative to ACC (left panel, blue bars, dashed line represents FDR<0.05) and pathways upregulated in ACC relative to HMSC (right panel, red bars, dashed line represents FDR<0.05) before filtering out cycling cells.
F. Distribution of cycling cells used to set a threshold for filtering (>0.3 cell cycling score) with ACC cells shown in red and HMSC cells shown in green.
G. UMAP clustering of all cells with cell cycling enrichment before (left) and after (right) filtering out cycling cells from (F).

**Supplementary Figure 2**

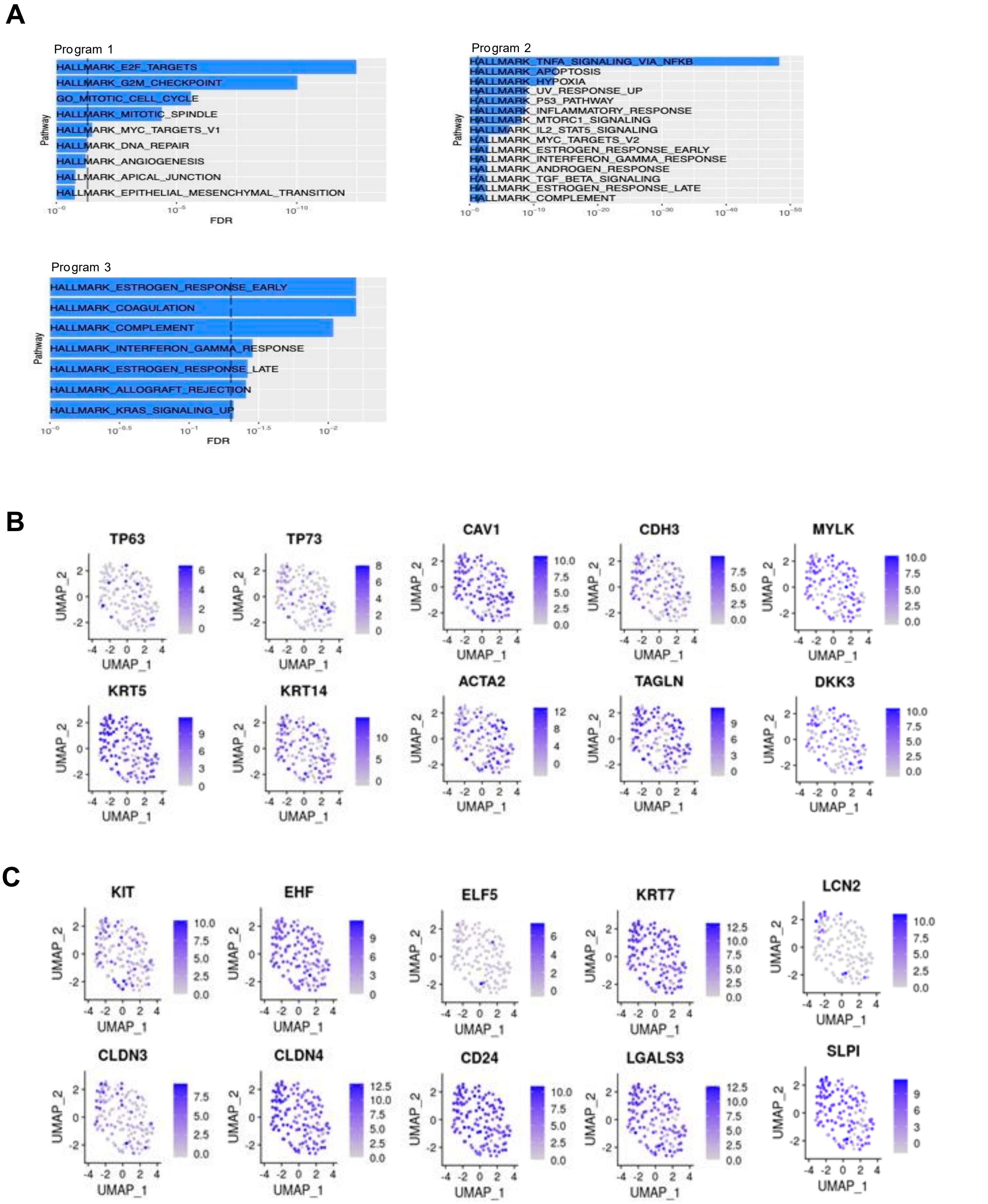

**Supplementary Figure 2. Additional analysis of myoepithelial and luminal markers in HMSC. Related to Figure 3.**

A. Bar plots show GSEAs of the three programs in (A) (dotted lines mark FDR<0.05).
B. UMAPs show expression for ten individual myoepithelial markers in HMSC cells, color indicates expression level.
C. UMAPs show expression for ten individual luminal markers in HMSC cells, color indicates expression level.

### Supplementary Figure 3

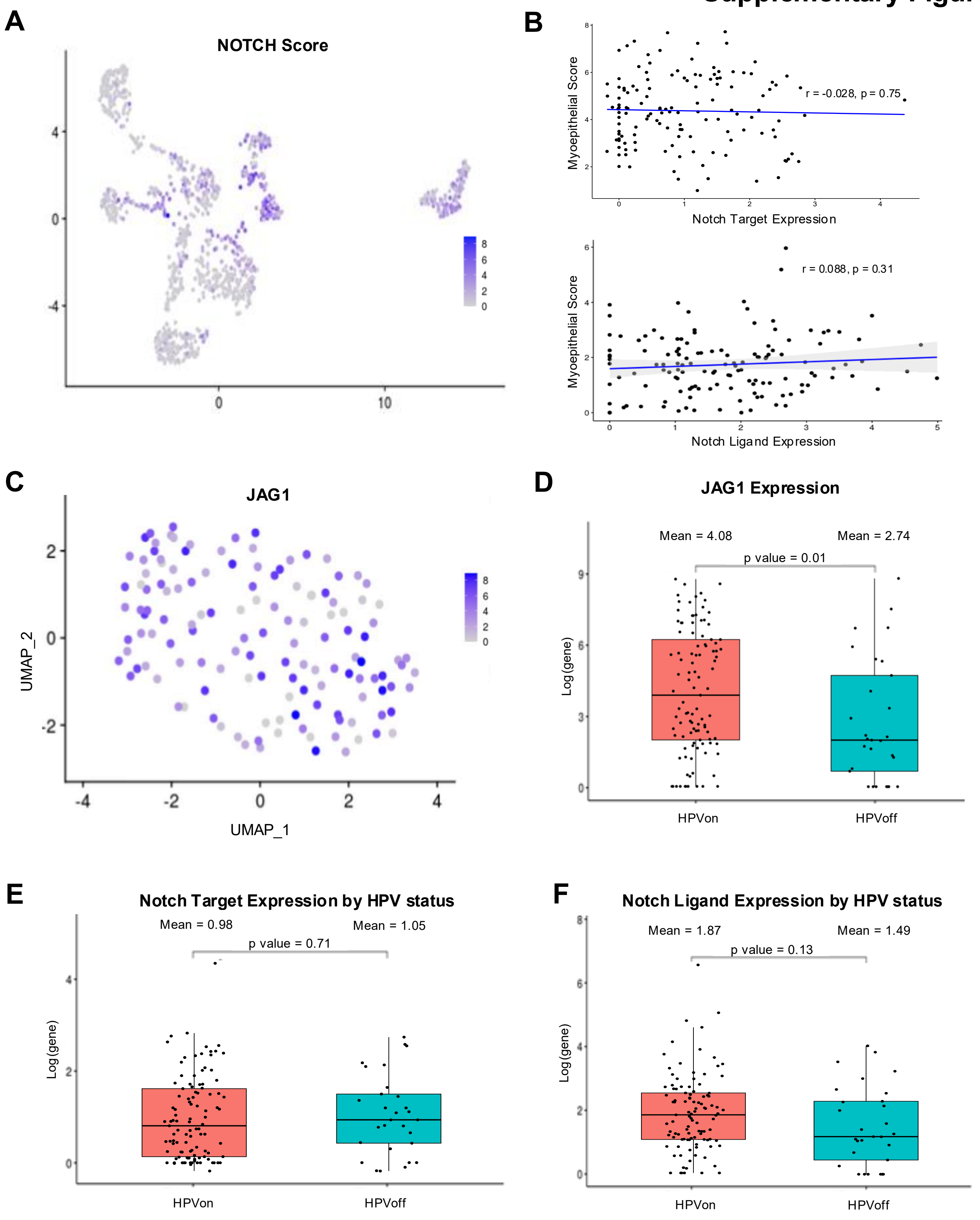

**Supplementary Figure 3. Additional JAG1/NOTCH expression levels. Related to Figure 4.**

A. UMAP clustering of HMSC and ACC malignant cells, colored by Notch expression score.
B. Scatterplots show non-significant correlation between Notch targets and myoepithelial score (top, p=0.75) as well as Notch ligands and myoepithelial score (bottom, p=0.31) in HMSC malignant cells without outliers.
C. UMAP clustering of HMSC malignant cells, colored by JAG1 expression level.
D. Boxplot shows higher JAG1 expression in HPVon cells (left) versus HPVoff cells (right) in HMSC (p=0.01).
E. Boxplot shows no difference in Notch target expression between HPVon cells (left) and HPVoff cells (right) in HMSC (p=0.71).
F. Boxplot shows no difference in Notch ligand expression between HPVon cells (left) and HPVoff cells (right) in HMSC (p=0.13).

#### Supplementary Figure 4

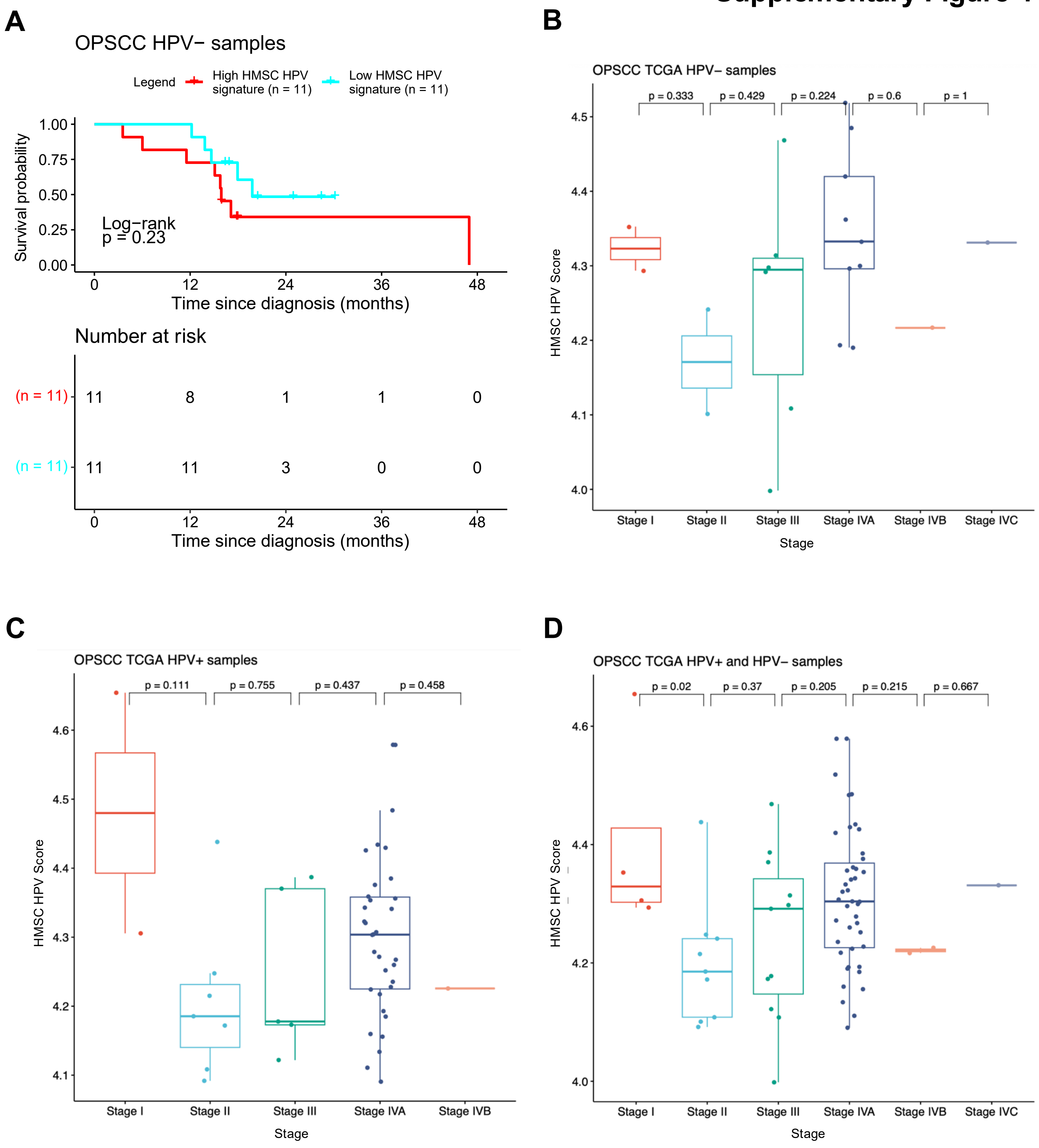

**Supplementary Figure 4. Additional survival analysis of HMSC HPV gene signature and tumor stage comparison in OPSC tumors. Related to Figure 6.**

A) Kaplan-Meier curves of TCGA HPV- OPSCC tumors stratified by expression of HMSC HPV signature. Red survival curves represent patients with high expression of HMSC HPV genes without a survival association (p=0.23).
B) Box plots of HMSC HPV signature in TCGA HPV- OPSCC tumors separated by stage.
C) Box plots of HMSC HPV signature in TCGA HPV+ OPSCC tumors separated by stage.
D) Box plots of HMSC HPV signature in TCGA HPV+ and HPV- OPSCC tumors separated by stage.

### Supplementary Figure 5

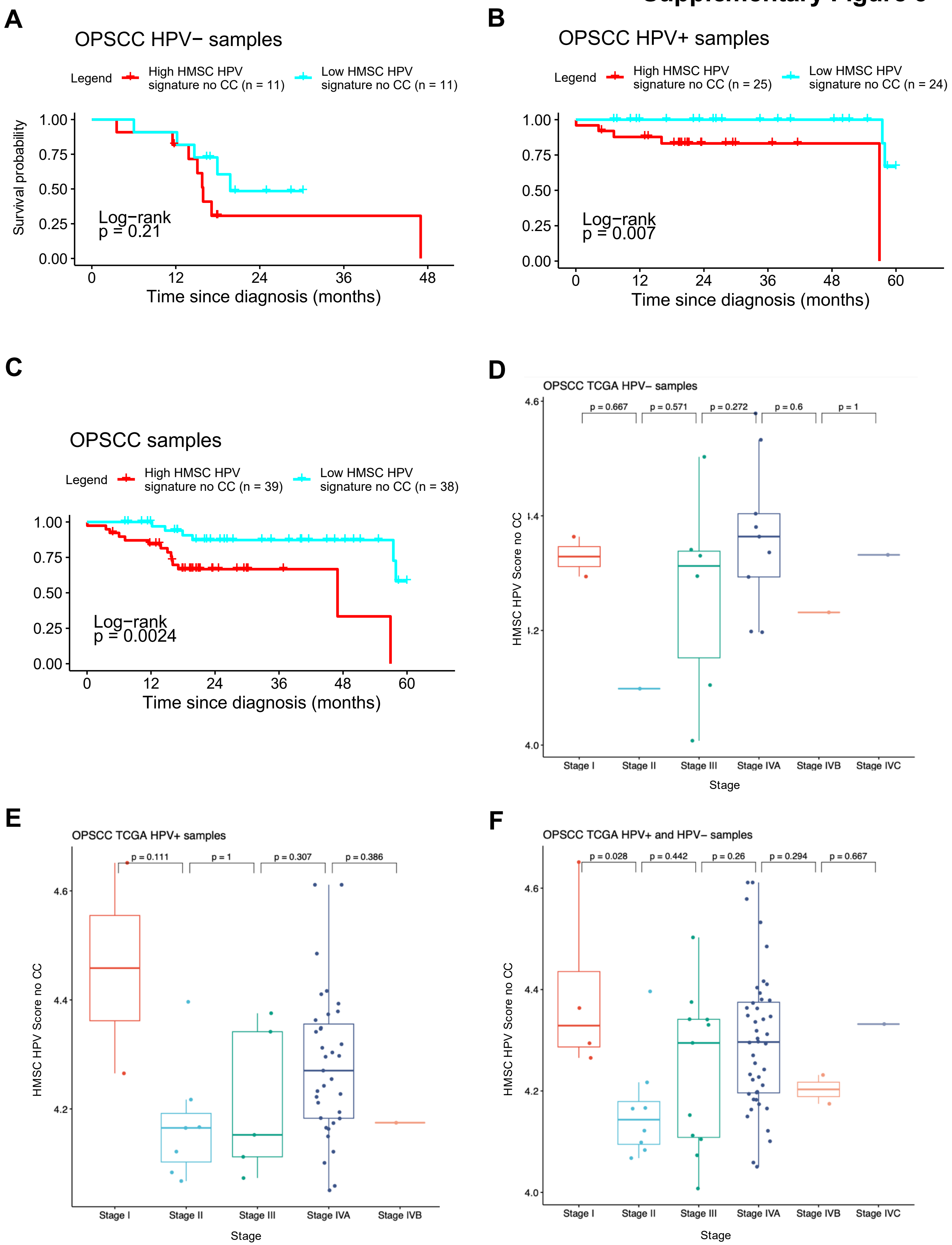

**Supplementary Figure 5. Additional survival analyses of HMSC HPV gene signature without cycling cells and tumor stage comparison in OPSC tumors. Related to Figure 6.**

A) Kaplan-Meier curves of TCGA HPV- OPSCC tumors after filtering out cell cycling genes stratified by expression of HMSC HPV signature. Red survival curves represent patients with high expression of HMSC HPV genes without a survival association (p=0.21).

B) Kaplan-Meier curves of TCGA HPV+ OPSCC tumors after filtering out cell cycle genes stratified by expression of HMSC HPV signature. Red survival curves represent patients with high expression of HMSC HPV genes and demonstrate poorer survival (p=0.007).

C) Kaplan-Meier curves of all TCGA OPSCC tumors after filtering out cell cycle genes stratified by expression of HMSC HPV signature. Red survival curves represent patients with high expression of HMSC HPV genes and demonstrate poorer survival (p=0.0024).

D) Box plots of HMSC HPV signature (after filtering out cell cycle genes) in TCGA HPV- OPSCC tumors separated by stage.

E) Box plots of HMSC HPV signature (after filtering out cell cycle genes) in TCGA HPV+ OPSCC tumors separated by stage.

F) Box plots of HMSC HPV signature (after filtering out cell cycle genes) in TCGA HPV+ and HPV- OPSCC tumors separated by stage.
